# Modeling unitary fields and the single-neuron contribution to local field potentials in the hippocampus

**DOI:** 10.1101/602953

**Authors:** Maria Teleńczuk, Bartosz Teleńczuk, Alain Destexhe

## Abstract

Synaptic currents represent a major contribution to the local field potential (LFP) in brain tissue, but the respective contribution of excitatory and inhibitory synapses is not known. Here, we provide estimates of this contribution by using computational models of hippocampal pyramidal neurons, constrained by in vitro recordings. We focus on the unitary LFP (uLFP) generated by single neurons in the CA3 region of the hippocampus. We first reproduce experimental results for hippocampal basket cells, and in particular how inhibitory uLFP are distributed within hippocampal layers. Next, we calculate the uLFP generated by pyramidal neurons, using morphologically-reconstructed CA3 pyramidal cells. The model shows that the excitatory uLFP is of small amplitude, smaller than inhibitory uLFPs. Indeed, when the two are simulated together, inhibitory uLFPs mask excitatory uLFPs, which might create the illusion that the inhibitory field is generated by pyramidal cells. These results provide an explanation for the observation that excitatory and inhibitory uLFPs are of the same polarity, in vivo and in vitro. These results also show that somatic inhibitory currents are large contributors of the LFP, which is important information to interpret this signal. Finally, the results of our model might form the basis of a simple method to compute the LFP, which could be applied to point neurons for each cell type, thus providing a simple biologically-grounded method to calculate LFPs from neural networks.

## Introduction

The local field potential (LFP) recorded from the hippocampus is rich in variety of signal during different network states. Sharp waves [9], ripples [11, 52], theta [8, 10], gamma [12] are different types of waveforms found in the LFP. These patterns of activity are population phenomena, which require synchronised contributions of large number of neurons. However, it was not until 2009 [16] and 2010 [2] that researchers showed that the LFP not only reflects synchronized network behavior, but also the field produced by just a single basket cell activity in the rat hippocampus *in vitro*. Previously field triggered by single neuron (called unitary field potential or uLFP) was thought to be of too small amplitude to be recordable above the noise level [39]. Why is hippocampal basket cell so special then? The axon of a basket cell does not extend very far from the cell body (soma) and it targets mostly the bodies and proximal dendrites of nearby pyramidal cells. In the hippocampus, pyramidal cell somata are packed in a single layer called stratum pyramidale, leading to the axon of a basket cell to form what appears to be the shape of a basket (hence the name). The synaptic currents induced in the postsynaptic population are therefore clustered in space.

However, in 2017 Telenczuk et al showed [44] that not only in the hippocampus but also in in the neocortex *in vivo* in human and in monkey, it is possible to extract unitary fields generated by not only single inhibitory but also by single excitatory neurons. Surprisingly, however, the two signals were of the same polarity despite being generated by currents of opposite sign. Moreover, there was a systematic time lag between them, with excitatory fields peaking later than inhibitory fields. It was hypothesised that excitatory uLFPs may be in fact di-synaptic inhibitory uLFPs: when a single pyramidal neuron fires, it induces the firing of inhibitory neurons which in turn generate the uLFPs. It is very likely that the same happens in the hippocampus where the pyramidal neuron-basket neuron connections are known to be very reliable [33].

In the present paper, we seek for plausible mechanisms to explain these observations, considering the hippocampus. We first reproduced the basket cell *in vitro* experiments in the model. We show that, indeed the extent of the axon of a basket cell creates high likelihood for triggering relatively large extracellular fields. We show how this signal spreads within different hippocampal layers. Next, we repeat the same simulations for two pyramidal cells with very different axon reach. Here, we show that the excitatory uLFP *in vitro* is of much smaller amplitude than the inhibitory uLFP, although the exact location and size will depend on the axon extent and where it is cut during the slicing procedure. Finally, we check if the hypothesis of Telenczuk et al [44] is also correct for the hippocampal data. By superimposing the excitatory uLFP with inhibitory uLFP after short delay we show that, indeed, the excitatory uLFP is being masked, leading to a pyramidal cell-triggered inhibitory field. Finally, we propose that uLFPs calculated by our model might form the basis of phenomenological models of the LFP, by convolving the generated spiking activity of point-neuron models with calculated unitary fields for specific cell types in space and time. This in turn will enable for better and faster understanding of recorded local field potentials.

## Materials and Methods

### Passive cellular models

Computational models were based on morphologically-reconstructed pyramidal neurons from rat hippocampal CA3 area. The morphologies were obtained from the *NeuroMorpo*.*org* online database and were integrated into the NEURON simulator [21] (Neuron 7.3) for simulations of the postsynaptic neurons. The NeuronEAP python library [43] (under Python 2.7) was used to calculate the local field potential. The time step of all the simulations was 0.025 ms. Passive membrane parameters were membrane resitance of *R*_*m*_ = 10000 Ω*cm*^2^, axial resistivity of *r*_*a*_ = 35.4 Ω*cm* and specific membrane capacitance of *c*_*m*_ = 1 *µf/cm*^2^. Other details about morphological arrangements are given in the Results section.

### Size of the slice

The soma of the presynaptic cell was assumed to be at coordinate (0,0,0). This assumption was used to calculate the time for the synapse onset (synapse placed further from the presynaptic cell soma would activate slightly later). The propagation velocity in the axon for inhibitory and excitatory neurons which we used for those calculations are indicated in the 1. We did not model the activity of the presynaptic cell. The slice size extended from −500 to 500 *µm* in length, −500 to 800 *µm* in height, and −200 to 200 *µm* (400 *µm*) in thickness as commonly used slice width in experimental studies interested in measuring local field potential [2, 29]. The somata of postsynaptic cells were placed throughout the length and the width of the slice and within −40 to 40 *µm* in height direction (ie. pyramidal cell layer).

### Postsynaptic population

To model the postsynaptic population, we inspected multiple CA3 pyramidal cell morphologies which were reconstructed from the rat hippocampus and which we downloaded from *neurmorpho*.*org* online database. This inspection was done in two ways: (i) visually, where we checked if the neurons did not look flatten and if the overall dendritic tree appeared uninjured (Fig. 1A), and (ii) quantitatively, where we monitored the change of size in diameter of the dendrites making sure that it decreased with the distance from the soma (Fig. 1B) as the diameter of the dendrites is of crucial importance for calculating the correct extracellular field. We decided to take all the selected reconstructions from the database of a single lab, which we chose to be the one of Amaral [24]. This selection process led us to 20 distinct CA3 pyramidal cell morphologies which we then translated vertically, with apical dendrites facing up (Fig. 1A). We randomly drew the morphologies from the pool of those 20 preselected cells to form the postsynaptic population. The number of segments varied between the cells but it was of scale of around 2000 segments per cell. The morphologies remained passive throughout the simulations. We decided to use only morphologies of pyramidal neurons, as they form the largest postsynaptic population, while other connections are mostly made to CA1 neurons [15, 27, 50, 6].

**Figure 1.**
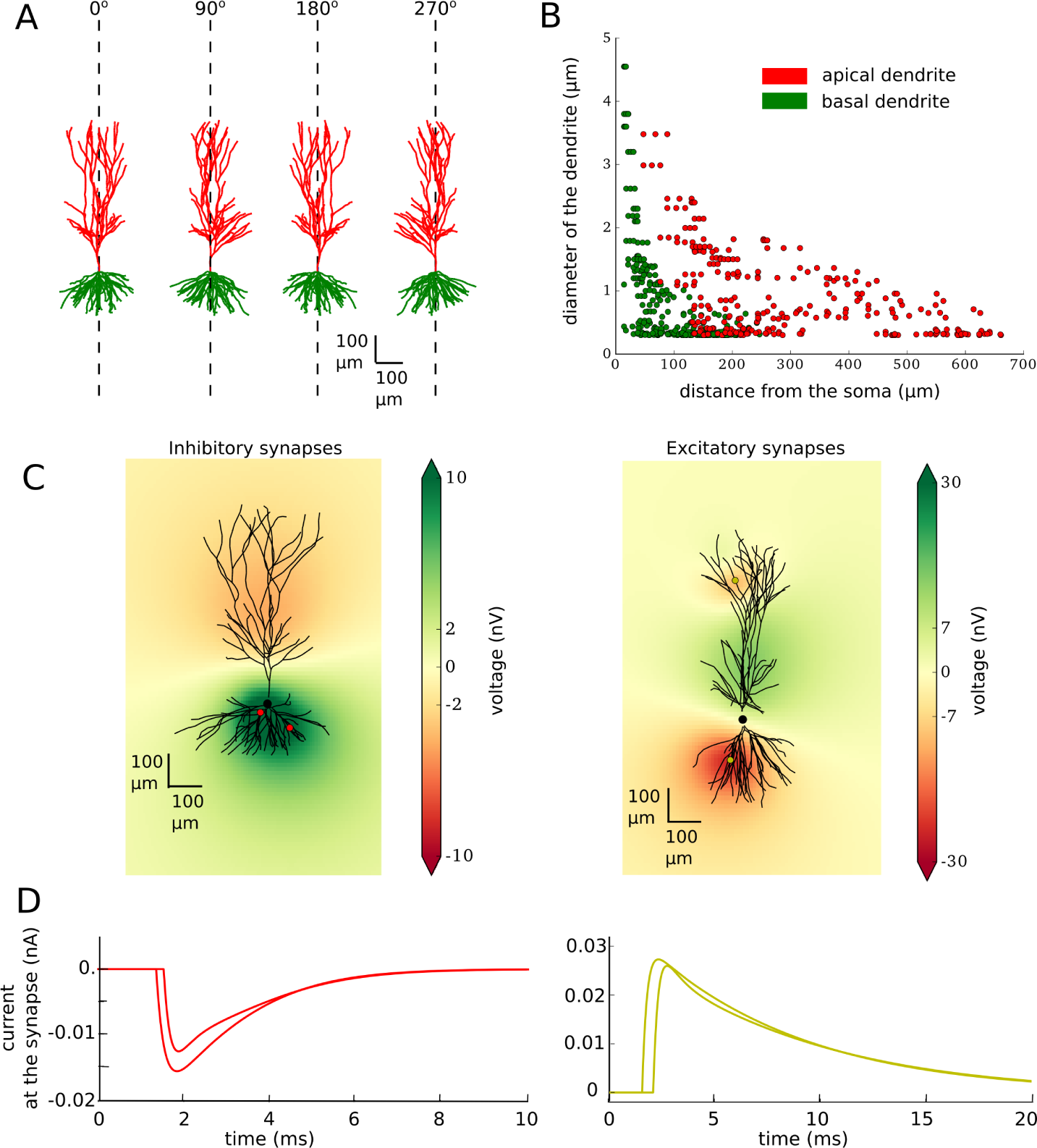
Model characteristics. **A**. Example of morphology used in the modelled population (20 different morphologies are used). All of the neurons are reconstructed uploaded by Amaral and can be downloaded from the *neuromorpho*.*org* (ID of the neuron shown in this figure: c81463). They are recorded in the rat CA3 area of the hippocampus. All of the neurons were translated to be vertically oriented with apical dendrites on the top and basal dendrites on the bottom. **B**. Width of the apical (red) and basal (green) dendrites as the function of their distance from the soma. **C**. Single neuron with two inhibitory (left) or two excitatory (right) synapses. Synapses are visualised as red (inhibitory) or yellow (excitatory) dots on the dendritic tree. Local field potentials is shown at 2.5 ms after beginning of the simulation (synapses were activated at 1 ms). **D**. Current at the inhibitory (left, red) and excitatory (right, yellow) synapses.

### Synaptic input

Next, we placed synapses on each of the postsynaptic neuron. Each synapse was placed directly on the dendrite. Parameters and number of the synapses (Table 1) differed for the two presynaptic cell types and were in the agreement with the literature. All postsynaptic neurons received at least one synapse; further synapses were added with probabilities indicated in the Table 1.

**Table 1.**
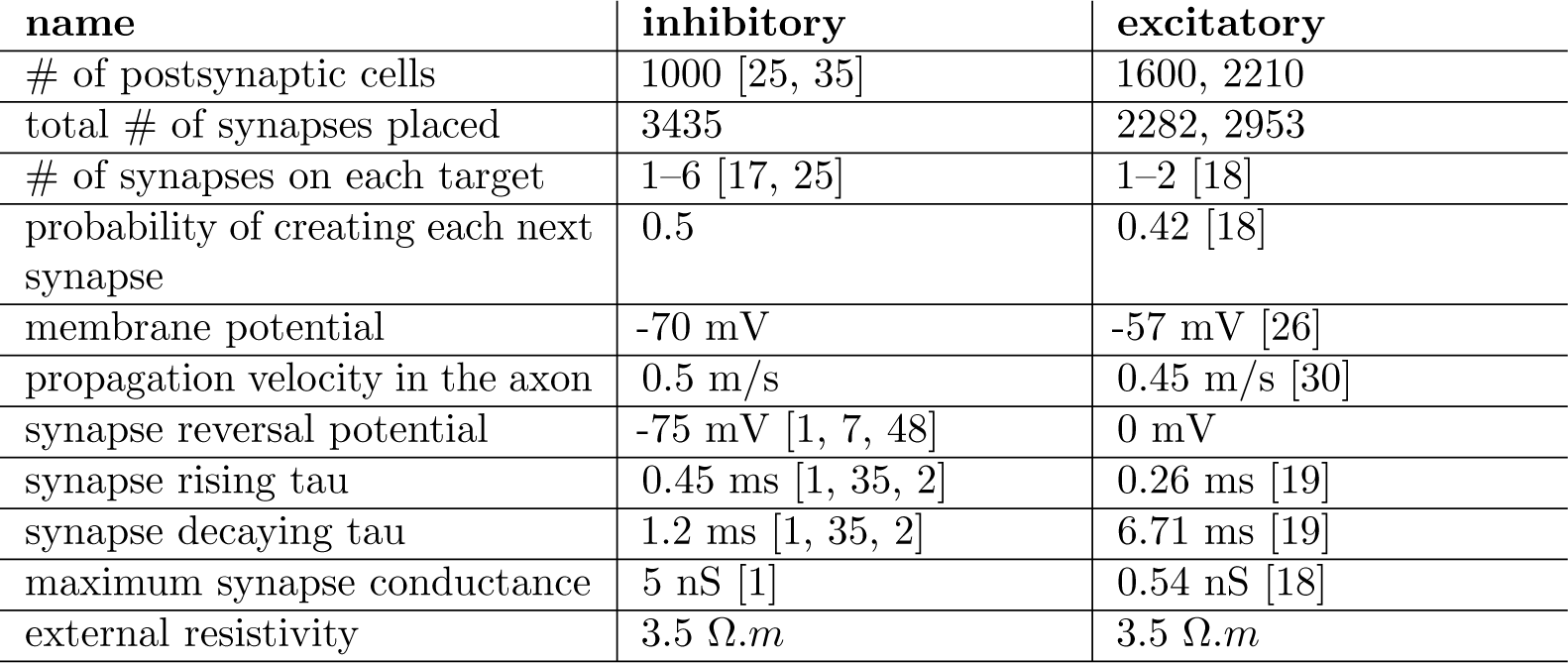
Cell parameters. Parameters used for modelling of basket cell (inhibitory) and pyramidal neuron (excitatory) together with the references to the original measurements. If under ‘excitatory’ column there are two different numbers given, first number is used for Cell A and the second for Cell B

| name | inhibitory | excitatory |
| --- | --- | --- |
| # of postsynaptic cells | 1000 [25, 35] | 1600, 2210 |
| total # of synapses placed | 3435 | 2282, 2953 |
| # of synapses on each target | 1–6 [17, 25] | 1–2 [18] |
| probability of creating each next synapse | 0.5 | 0.42 [18] |
| membrane potential | -70 mV | -57 mV [26] |
| propagation velocity in the axon | 0.5 m/s | 0.45 m/s [30] |
| synapse reversal potential | -75 mV [1, 7, 48] | 0 mV |
| synapse rising tau | 0.45 ms [1, 35, 2] | 0.26 ms [19] |
| synapse decaying tau | 1.2 ms [1, 35, 2] | 6.71 ms [19] |
| maximum synapse conductance | 5 nS [1] | 0.54 nS [18] |
| external resistivity | 3.5 $\Omega.m$ | 3.5 $\Omega.m$ |

The amplitudes and time constants of simulated synaptic currents are also given in Table 1. Synaptic current is usually measured from the soma, which is not a problem in case of the basket cells, which place their synapses in the soma. However, it may cause discrepancies in case of input from pyramidal neurons which place their synapses far from the soma. To account for this we used the values calculated for the current at the dendrite as given in the paper of Guzman and colleagues [18].

### Calculation of the local field potential

To calculate local field potential generated by activation of the synapses on each neuron in space, we used the NeuronEAP python library [43] which is based on the linear source approximation which calculates the summed potential generated by currents originating from line sources with known sizes and positions [22, 49]. In all calculations, we used an extracellular conductivity of 0.3 Sm [38]. Figure 1C shows an example of local field potential for two randomly placed inhibitory (left) and excitatory (right) synapses. The current at each of the synapse is plotted in Fig. 1D. Note the difference in latency caused by the axonal propagation delays.

### Active cellular models

In some control simulations, we used voltage-dependent channels, which were taken from models of hippocampal pyramidal cells developed previously [32, 46]. The active cell models had voltage-dependent *Na*^+^, *K*^+^ and h-type channels distributed through the cell, with densities of 5 mS/cm^2^ for Na^+^, 5 mS/cm^2^ for K^+^ and from 5 to 10 mS/cm^2^ for *I*_*h*_. In this configuration, the LFP generated by excitatory synaptic inputs was affected by the presence of voltage-dependent currents (Fig. 2, top panels), diminishing the LFP amplitude, with a peak effect close to 1 *µ*V, and generally smaller. In contrast, the LFP from inhibitory inputs was little affected by the presence of voltage-dependent currents (Fig. 2, bottom panels). Because of these limited effects, and the fact that the simulation time is considerably larger with voltage-dependent currents, we considered only passive neurons in simulations involving large populations of morphologically-reconstructed neurons.

**Figure 2.**
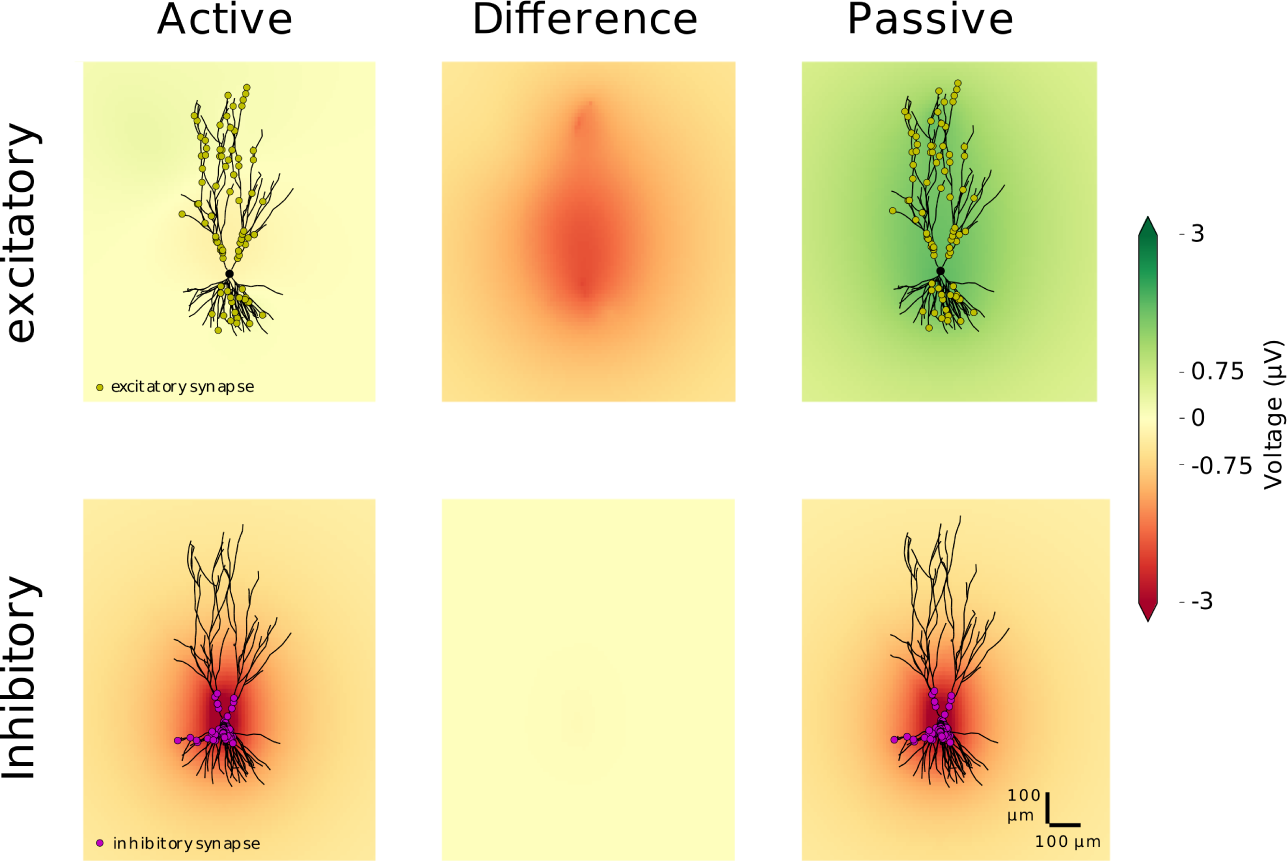
Impact of voltage-dependent conductances on single-cell LFPs. The LFP was calculated from the activation of synapses in single-cell simulations, comparing passive and active neurons. The active cell had additional *Na*^+^, *K*^+^ and h-type channels distributed through the cell (see Methods). 100 synapses, excitatory (top) or inhibitory (bottom), were placed on a single postsynaptic pyramidal cell (NeuroMorpho.Org ID: NMO 00199 [24]) according to their biological localisation (excitatory synapses distanced from the soma and inhibitory synapses at the soma and nearby dendrites). The colored field is the field generated by the activation of the synapses on the neuron with active channels (left) and on the passive neuron (right). The field is displayed at the time point with the largest absolute difference between the field produced by the active and passive neuron. The difference at this time point is shown in the middle panels.

### Code accessibility

The code for the figures of the full morphology model is deposited and available at Zenodo [45]. To calculate the local field potential, we used the NeuronEAP Python library [43].

## Results

### Inhibitory unitary field potential

First, we reproduced published experimental results of Bazelot and colleagues [2] in the model. We placed 1000 pyramidal cells in space to mimick the slice configuration [35] (as indicated in Materials and Methods; location of the somata of the postsynaptic cells: Fig 3A). Next we created at least one and maximum of six inhibitory synapses on each of the cells. The highest probability of creating a synapse was within the pyramidal cell layer or, within stratum lucidum [35]. Throughout the length of the slice the probability decreased with the distance from the body of the presynaptic cell with a gaussian profile. The exact location of the synapses is indicated by red dots in Fig. 3B. Red histograms show the distribution of the synapses throughout the length of the slice (Fig 3B top histogram) and throughout the hippocampal layers (Fig 3B histogram on the right). Four randomly chosen morphologies of postsynaptic neurons with somata represented by black dots were also drawn to give an idea of the spread of dendritic trees through the layers (Fig 3B).

**Figure 3.**
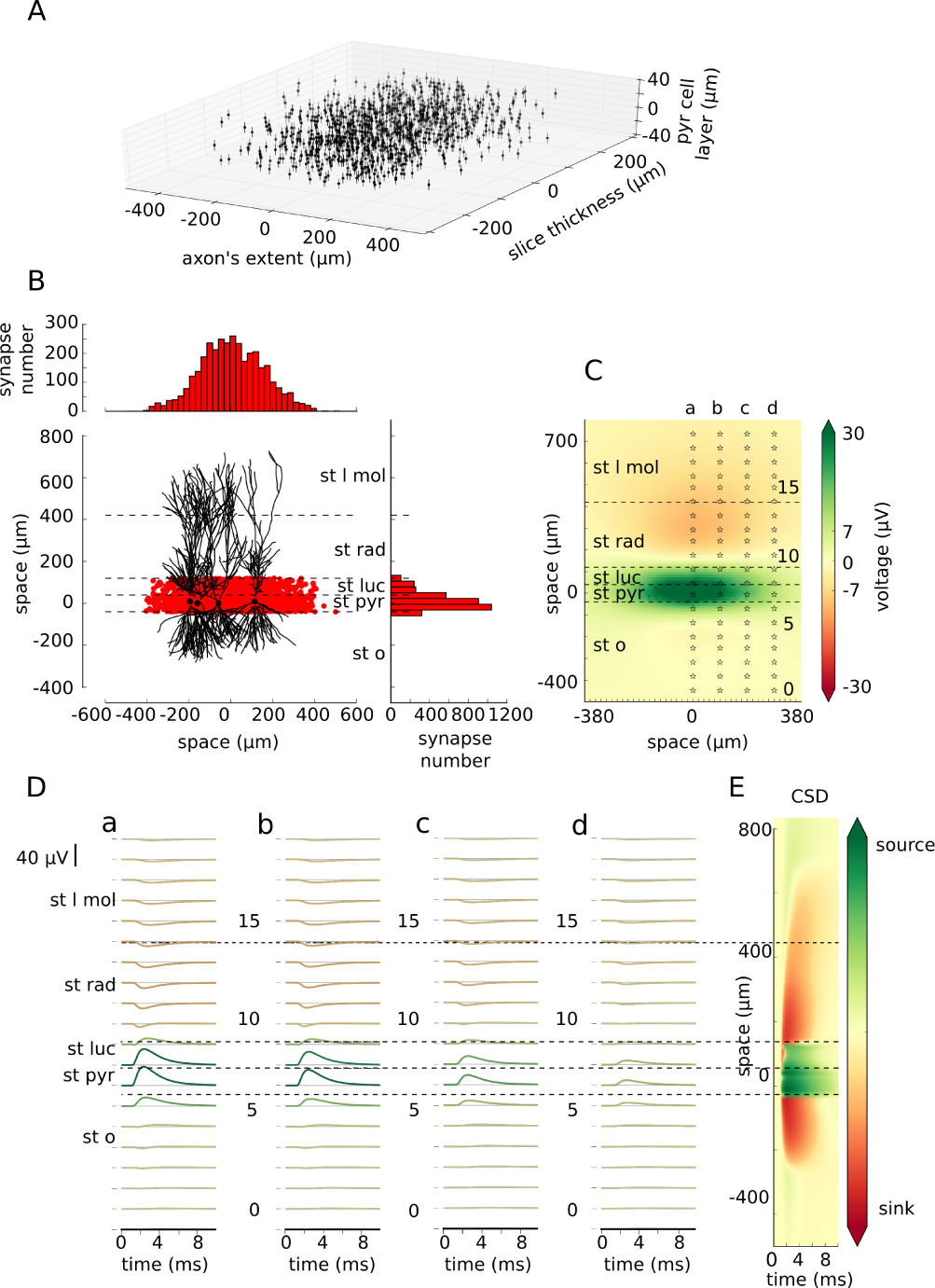
Inhibitory unitary field potential. **A**. Distribution of the postsynaptic neurons within a slice. Each dot represents a soma of one pyramidal cell. **B**. Distribution of the synapses. Each red dot shows the location of the inhibitory synapse within the length of the slice and within the hippocampal layers. 3435 inhibitory synapses were placed on the postsynaptic targets. Their distribution in both axes are shown on the top and on the right. Four, randomly selected, exemplary postsynaptic neurons with their somata indicated by black dots are drawn for better understanding of spatial relations. **C**. Local field potential at 2.5 ms from the start of the simulation (synapses were activated at 1 ms). Stars show the locations of the electrodes (0–19) with each fifth electrode marked by a number. Electrodes form 20-electrode arrays marked a–d. **D**. Traces recorded by the electrode array a–d corresponding to the locations from C. Traces are coloured by their maximum absolute peak corresponding to the colormap in (C). E. Current source density analysis done on the field average across the length of the axon (x direction). Layers: st l mol – *stratum lacunosum moleculare*, st rad – *stratum radiatum*, st luc – *stratum lucidum*, st pyr – *stratum pyramidale*, st o – *stratum oriens*.

Next, we simulated the activation of the synapses and we calculated how the generated current spreads through the cells and in the extracellular space. From those currents we calculated the LFP within 10 ms of the simulation time. An example of LFP (1.5 ms after the activation of the closest synapses) is shown in Fig. 3C. The LFP is shown across different layers of the hippocampus: stratum lacunosum moleculare (St l mol), stratum radiatum (st rad), stratum lucidum (st luc), stratum pyramidale (st pyr) and stratum oriens (st o). Not surprisingly, the potential of the highest amplitude is recorded around the location of the synapses. Columns of stars marked a–d represent the location of the array of electrodes placed along the hippocampal layers. Each electrode in an array is numbered 0-19. Such recordings of local field potential in the CA3 area of the hippocampus *in vitro* have been previously performed experimentally using 8 electrodes [2, 3]. The traces obtained from each electrode are shown in Fig. 3D. Their amplitude decreases with the distance of the presynaptic neuron with agreement to Bazelot et al. (2010) [2]. The location of the electrode has an influence on the amplitude and deflection of the recorded signal. Finally, we calculated current source density analysis which clearly shows the source of the current in the pyramidal cell layer and nearby.

Next, to compare our findings with the published experimental results we selected one of the largest signals (array a, electrode 7) and we measured its amplitude, and time from the beginning of the rise to the peak of the signal (Fig 4A). In the paper of Bazelot and colleagues [2] the mean amplitude of the recorded signal was 28.1 *µV* whereas recording from our largest waves was 36.7 *µV*. Although, Bazelot and colleagues did not specify rise-to-peak time, the timings read from their figures are similar (1.53 ms in the Fig. 4A). After that, we checked how the location of the maximum and minimum peak of the signal varies depending on the location of the electrode in different layers. To this extent we took measurements from all the electrodes in the electrode array and we checked for the maximum and minimum in time. The time of the peaks varied largely depending where the electrode was placed (Fig 4B). Finally we measured the peak to peak deflection throughout different layers, distribution of which we show in Fig 4C. It shows how the amplitude and the deflection of the measured signal might change with just a very slight shift of the electrode within the hippocampal layers.

**Figure 4.**
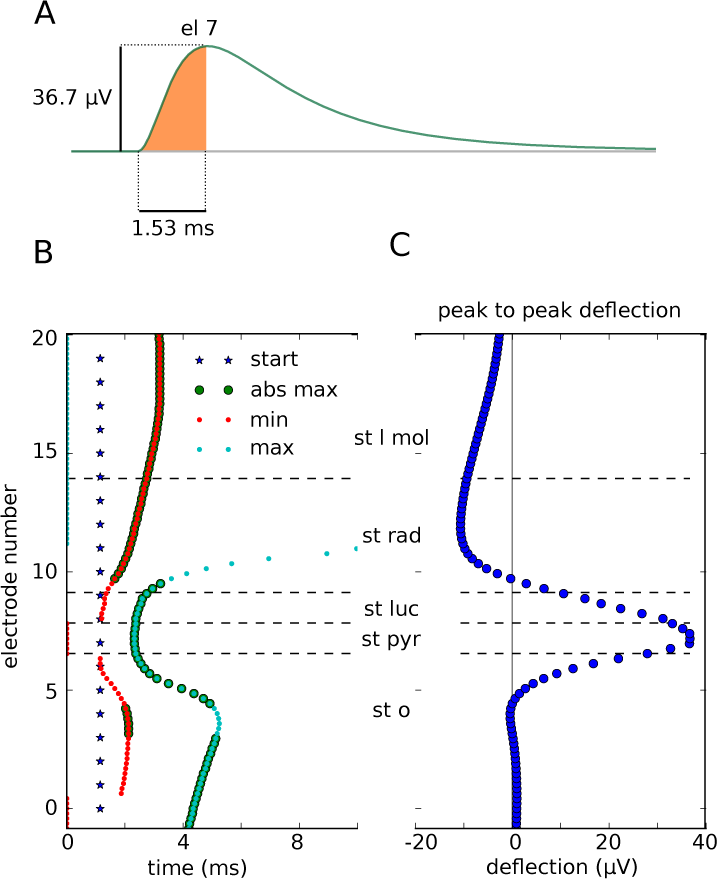
Characteristics of inhibitory uLFPs. **A**. Recording from the electrode 7, array a (location shown in Fig 3C). The area shaded in orange indicates the measurements: the amplitude of 36.7 *µV* and time to peak of 1.15 ms. **B**. Stars show the beginning of the synapse activation. Time to minimum and maximum peak of each trace recorded by the array a is indicated by red and blue dots respectively. Enlarged dots indicate if the peak was absolute maximum in the trace. Time to peak varies between layers. The peak arrives the earliest (start of the rise to peak: 1.53 ms) in stratum pyramidale while it is as late as 3.05 ms in stratum moleculare (time from blue line to bold red and blue dots). **C**. Peak to peak deflection within different hippocampal layers. The highest positive peak is in stratum pyramidale but it points downwards in in stratum radiatum and stratum lacunosum moleculare

### Excitatory unitary field potential

Axonal trees of pyramidal neurons are very different from those of basket cells. They tend to be very long (200 mm for CA3b to 500 mm for CA3c pyramidal neuron [40], as compared to 900–1300 *µm* in basket cells [25]) and longitudinal projections of single axons can extend very far (even 70% of the dorso-ventral extent of the hippocampus) [27, 28, 42] (but see [37] in Dentate Gyrus).

To model uLFPs produced by single pyramidal neurons, we used morphologically-reconstructred pyramidal neurons from rat CA3 (see Methods). We searched the NeuroMorpo.org online database for best preserved pyramidal cell axons from the rat CA3. We selected two cells: one with *NeuroMorpho*.*Org* ID: NMO 00187 [47] which we will call Cell A (Fig. 5A left) and second cell with *NeuroMorpho*.*Org* ID: NMO 00931 [41], which we will call Cell B (Fig. 5A right). Next, we rotated them so that the dendritic tree was oriented vertically and we calculated the length of the axon in each 50*µm* x 50*µm* bins. Blue histograms in Fig. 5A left and right show the total length of the axon within 50 *µm* bin in each axis (and summed across other axes) for Cell A and Cell B respectively (the length of the axon in the z-direction is not shown). Axon is drawn in blue and the location of the soma is indicated by the red star. Next, we cut the axon to the slice of size: −500 *µm* to 500 *µm* from the soma of the presynaptic pyramidal cell in x-direction and by −500*µm* to 800*µm* in y-direction and by −200*µm* to 200*µm* in z-direction. The extent of the slice in two directions is shown by green rectangle and the remaining length of the axon by green histograms in Fig. 5A. We calculated total length of the axons by adding all the measurements from all 202 the bins. Total length of the axon of Cell A was 468.57 mm, after the cutting only 11.16 mm remained (being around 2% of the original axon). Total length of the axon of Cell B was 205.17 mm, after cutting 14.12 mm remained (around 7% of the original axon). By giving this numbers we want to emphasize how little fraction of the pyramidal cell axon remains in the experimental slice. This has been also pointed out previously [23].

**Figure 5.**
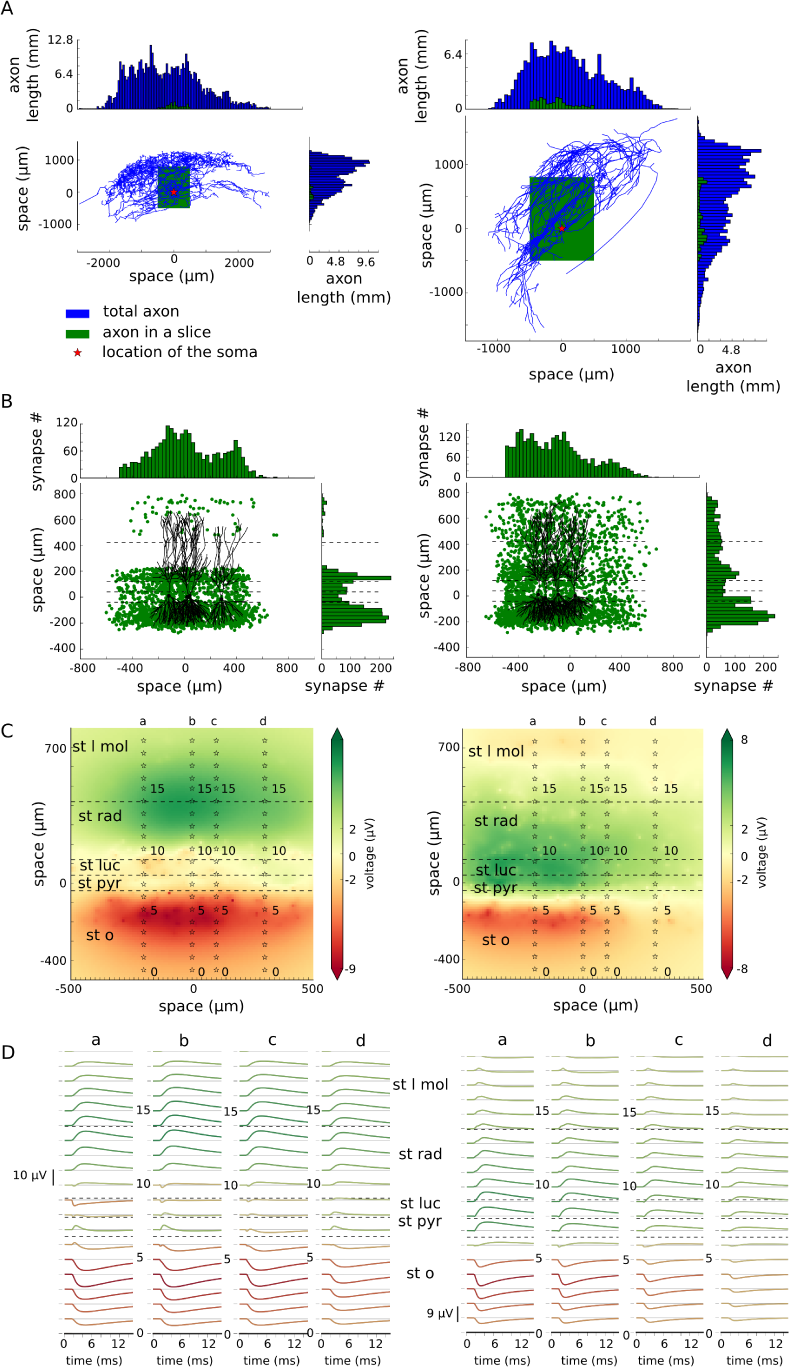
Excitatory unitary field. **A**. Axon morphologies of two CA3 pyramidal cells, Cell A (left, ID: NMO 00187) and Cell B (ID: NMO 00931) downloaded from *neuromorpho.org*: These cells were rotated so that their dendrites are placed vertically. Blue histograms show the length of the axon in each 50 *µm* bin in two axes. Axon morphologies are indicated by blue lines with the red star showing the location of the soma. Green rectangles shows where the axon was cut consistently with the size of a typical slice (−500 *µm* to 500 *µm* from the soma in the length of the slice, −500 *µm* to 800 *µm* in the height and −200 *µm* to 200 *µm* in the width of the slice). Green histograms show the length of the axon remaining after the cutting. **B**. The distribution of the excitatory synapses in the model for Cell A (left) and Cell B (right). The distribution follows the distributions calculated by the length of the axon in A, however with constraints given by the morphologies of the postsynaptic cell population. Four randomly chosen morphologies of postsynaptic cells were drawn for easier visualisation of the spatial relations. **C**. Local field potential plotted at 5.5 ms after beginning of the simulation with four electrode arrays (a–d) placed at −200, 0, 100 and 300 *µm* away from the presynaptic cell soma. **D**. Traces recorded by each of the electrode arrays marked as a–d.

It is known that inter-varicosities distance on the CA3 pyramidal axon is on average 4.7 *µm* [42, 27, 51]. We combined this information with the calculated length of the axon to estimate the probability of placing a synapse within each 50*µm* bin. Total number of synapses placed by Cell A should be around 2400 and placed by Cell B it should be around 3000. CA3 pyramidal neuron in 58% cases places 1 synapse on its postsynaptic target and in remaining 42% cases it places 2 synapses [19]. Therefore we created the postsynaptic cell population of cell A to be 1600 and of cell B to be 2210 cells. We gave the probability of placing a synapse matching the distribution of the cut axon, by doing so we ended up with the synapse distribution as indicated by green dots and green histograms in Fig. 5B (left and right for Cell A and B respectively). Cell A placed 2282 synapses and Cell B placed 2953 synapses on its postsynaptic targets.

Next, we calculated local field potential generated by the two neurons. The snapshot of those calculations at time 5.5 ms from the beginning of the simulation is depicted in Fig. 5C (Cell A, left; and Cell B, right). Here, we placed 4 electrode arrays at −200 *µm*, 0 *µm*, 100 *µm* and 300 *µm* from the presynaptic cell body (stars in Fig. 5C indicated by a–d) because due to the non-symmetric axon, the synapse distribution is also non-symmetric. Unitary field potentials recorded by each of the electrodes (Fig 5D) differ largely from those recorded by activation of basket cell synapses. As expected, the distribution of uLFPs depends on the shape and the extent of the axon. uLFPs for cell A reaches amplitude no larger than 10 *µV*. However, at st pyramidale where recordings are most frequently performed the uLFPs are of amplitude near 0 *µV* (peak-to-peak in electrode 7 array b is 2.2 *µV*), with the highest amplitude (up to 8 *µV* in peak to peak measurements in the array b, and 8.53 *µV* in el 4, array a) in the distant layers such as stratum radiatum and stratum oriens (Fig 5D. The uLFP generated by cell B is of different distribution. Here, the signal is comparably large in the pyramidal cell layer, even though the strongest signal can be found in st oriens and towards the left part of the slice (Fig 5D a right, 200 *µm* away from the soma of the presynaptic cell). However, even the highest uLFP is still of the amplitude not larger than 9 *µV*.

Finally, we checked the timing of the highest and lowest uLFP peaks within different layers of the hippocampus at location 0 *µm* from the presynaptic cell body (Fig 6A). The time of the absolute maximum peaks differed by as much as 6 ms depending on the location of the measurement. The profile of the peak-to-peak deflection differed between the two cells (Fig 6B left, Cell A; right Cell B) and it changed across the different layers.

**Figure 6.**
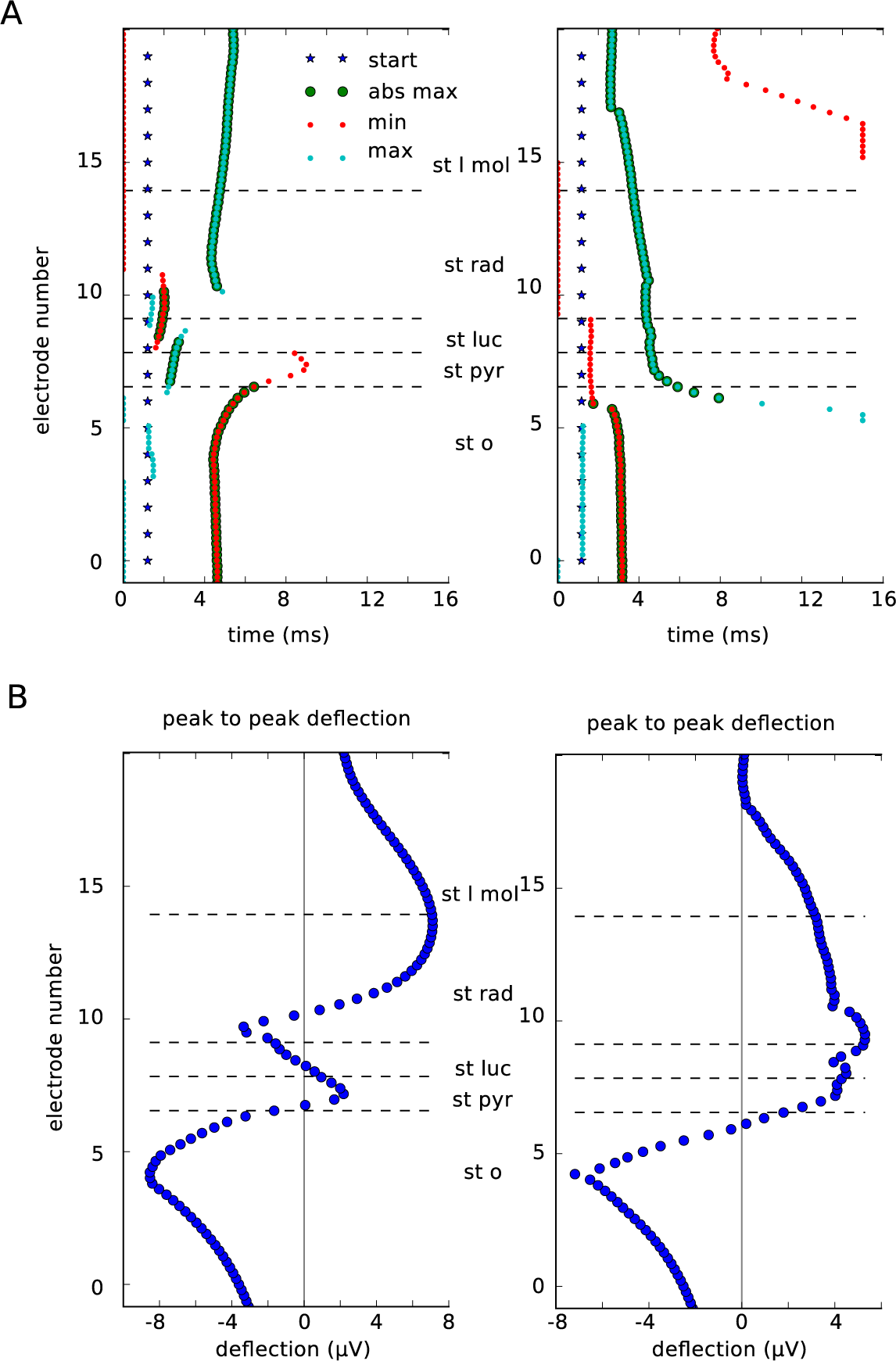
Characteristics of excitatory uLFP. Results for Cell A are on the left and for Cell B are on the right. **A**. Stars show the beginning of the synapse activation. Time to minimum and maximum peak of each trace recorded by the array b in Fig 5C (0 *µm* from the presynaptic soma) is indicated by red and blue dots respectively. Enlarged dots indicate if the peak was absolute maximum in the trace. Time to peak vary between layers. **B**. Peak to peak deflection within different hippocampal layers. It varies between the two cells (left and right).

We conclude that for the two pyramidal neurons the excitatory uLFP might prove difficult to measure experimentally *in vitro*. One would need to place an extracellular electrode in the correct location which differs from cell to cell.

### Masking of excitatory uLFP with inhibitory uLFP

Pyramidal cells form only few synapses on their basket cell targets, however, those connections are known to be very reliable [33]. Recently, Telenczuk and colleagues proposed that the unitary fields triggered by the activation of the excitatory neurons which we recorded from the human and monkey neocortex were in-fact bi-synaptic inhibitory unitary fields [44]. We believe that this might also be true in the hippocampus. To check if this is indeed plausible, we superimposed the excitatory uLFPs generated by Cell A and Cell B with the inhibitory uLFP after a 3 ms time delay (Fig. 7) [33, 34]. The local field potential at 5.5 ms after the beginning of the simulations shows much stronger contribution of the inhibitory uLFP with very strong positive field around stratum pyramidale (Fig 7A). The recordings from the a–d electrode arrays reveal very minor excitatory uLFP contribution compared to the strong inhibitory uLFP contribution (Fig 7). Our results show that, indeed it might be difficult to separate excitatory uLFP from the inhibitory one without use of manipulations that would block specific cell types. Please also note that in our model both inhibitory and excitatory neurons are located at (0,0,0) coordinate therefore the signal is strong for both. However, in the real recordings it is more likely that the somata will be shifted.

**Figure 7.**
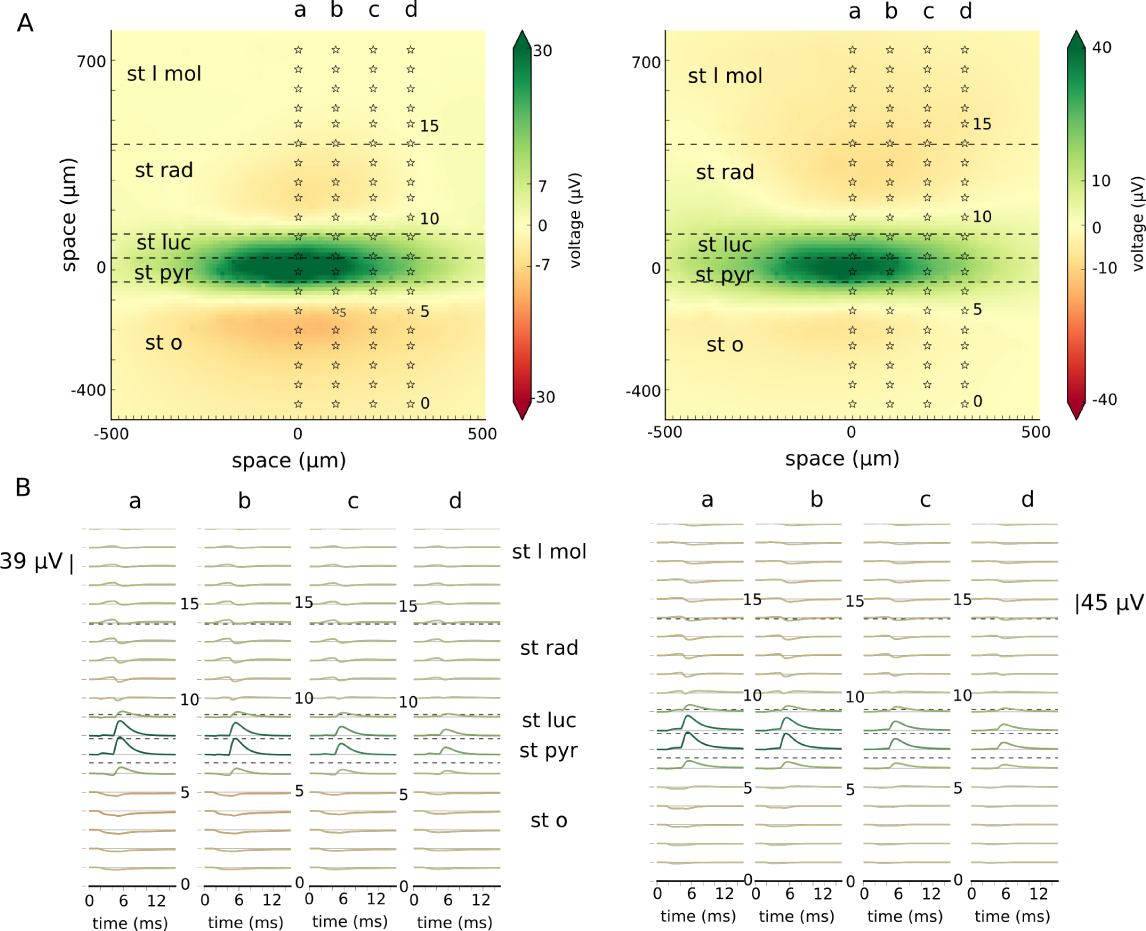
Masking of excitatory uLFP with inhibitory uLFP. Results for Cell A are on the left and for Cell B are on the right. **A**. Local field potential at time 5.5 ms from the beginning of the simulation. At time 1 ms excitatory synapses were activated (of Cell A on the left, of Cell B on the right) followed by the activation of inhibitory synapses at time 3 ms. Stars show the location of the electrodes belonging to the arrays marked a–d. **B**. Traces showing recordings from the electrode arrays marked a–d in A.

## Discussion

In this paper, we have used numerical simulations of morphologically-reconstructed neurons to investigate the single neuron contribution to local field potentials, the so-called unitary local field potentials or uLFPs. In agreement with previous studies in hippocampus [2, 16] and neocortex [44], we found that inhibitory uLFPs are of larger amplitude than excitatory uLFPs. Consequently, the LFP signal is expected to be dominated by inhibitory currents. We discuss below these findings, their significance and what perspectives they offer for further work.

Our biophysical model was based on reproducing published experimental results [2] of inhibitory unitary field in the hippocampal CA3 slice from the rat. Next, we used the same model to find out what is the excitatory unitary field produced by pyramidal neurons in the same area. We show that pyramidal neurons also produce unitary field potentials, however of much smaller amplitude and of very different spatial profile which depends on their exact axonal architecture. Due to limited computational resource constraints we were unable to calculate the field generated by the full pyramidal cell axon (*in vivo* condition). However, if such resources are available, it would be of interest to check if the excitatory uLFP remains of the same amplitude if the whole axon morphology is considered.

By comparing the two types of uLFPs, we found that it is likely that the excitatory uLFP is further masked by the inhibitory uLFP triggered by pyramidal– basket cell interaction.

The explanation for the dominance of inhibitory uLFPs is based on the par-ticularities of the pyramidal cell morphology, as illustrated in Fig. 8. Excitatory synapses, which are located exclusively in apical, oblique and basal dendrites, produce single-synapse LFPs which are of various polarities, according to their positions [13, 17, 31]. For example, basal dendrite synapses and apical synapses will produce dipoles of opposite polarity, so will partially cancel (Fig. 8A). This cancellation explains why the uLFP of excitatory synapses is of relatively small amplitude. Inhibitory synapses on pyramidal cells also contact the various parts of the dendrites and will suffer from the same cancelling effect (Fig. 8B). This cancellation will thus also occur even if inhibitory synapses are depolarizing. However, inhibitory synapses have in addition a very high density in the perisomatic region, which not only causes strong inhibition, but it also always forms the same dipole. These dipoles on each pyramidal cell sum up, and yield a uLFP of larger amplitude (Fig. 8C). This explanation suggests that the spatial distribution of synapses in the cell, and its asymmetry, determine the respective excitatory and inhibitory contributions to LFPs. This explanation is supported by our computational models.

**Figure 8.**
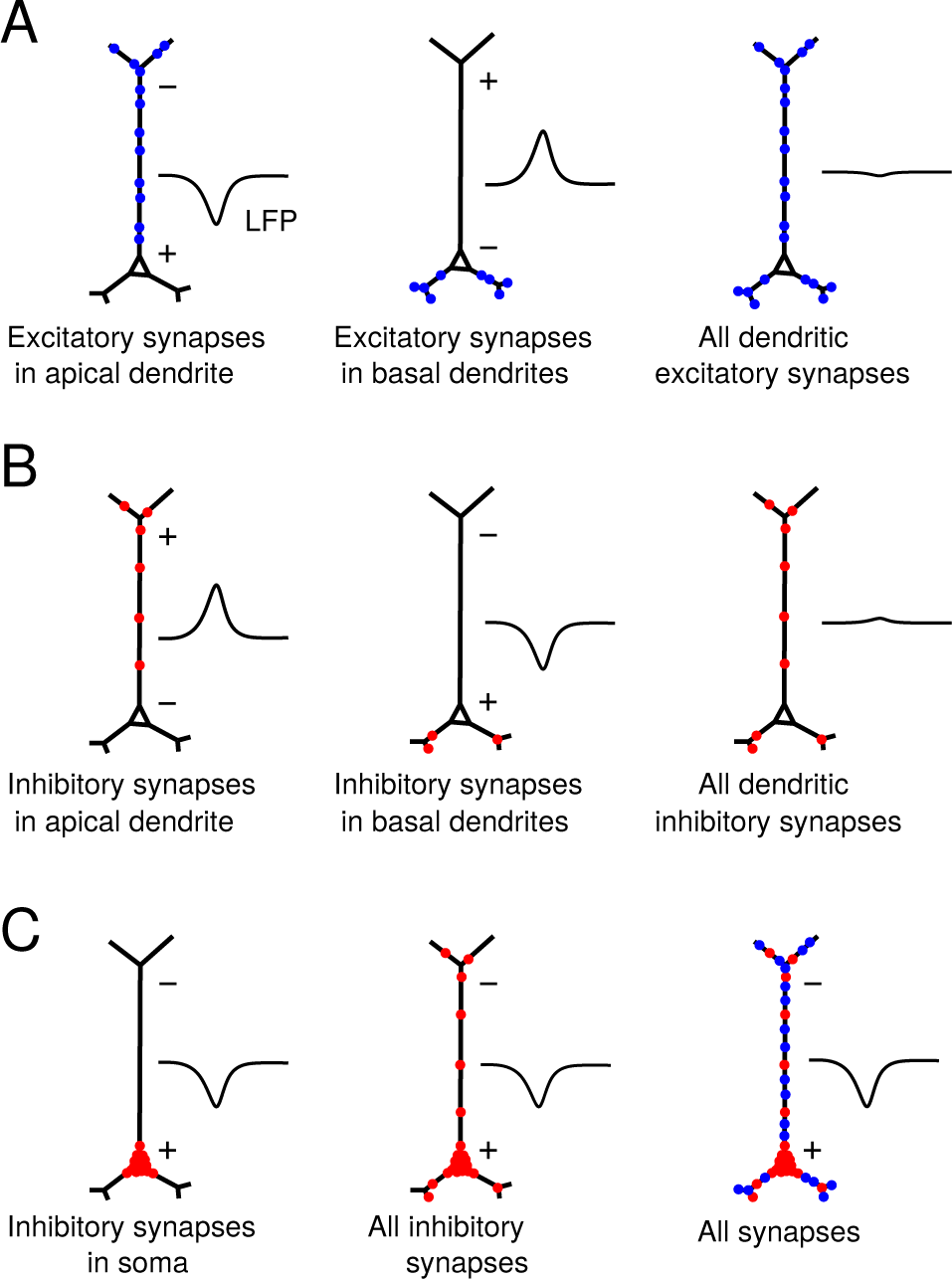
Proposed biophysical origin of the dominant contribution of somatic inhibition in the local field potential. **A**. Excitatory synapses occurring in the apical dendrite (left) or in basal dendrites (middle) produce dipoles of opposite sign. All dendritic synapses therefore produce a LFP of moderate amplitude (right). **B**. The same cancellation applies to inhibitory synapses in apical and basal dendrites. **C**. Inhibitory synapses in the soma always form the same dipole (left), which will dominate when all inhibitory synapses are present (middle) or with all synapses (right).

In addition, we showed that the axon morphology of pyramidal neurons has a critical influence on the uLFP recorded along the radial and lateral axes (Fig. 5). The morphologies of the axon can vary drastically across neurons, which in turn determines the final distribution of the synapses on their target cells. Importantly, most slice preparations cut significant part of the axonal arbor leading to a pronounced decrease in the number of synaptic terminals, which could additionally weaken the effect of pyramidal neurons on the LFP compared to the inhibitory neurons.

Inhibitory neurons are generally thought not to contribute to the LFP due to their spherical symmetry which generates a closed-field geometry that produces little electric field [28]. This argument holds mainly for the far-field potentials directly resulting from interneurons. Here, we show that inhibitory neurons contribute significantly to the LFP, through their postsynaptic effect on pyramidal cells. Thus, we consider here the post-synaptic contribution of the neurons, which does not depend on the dendritic shape of the pre-synaptic inhibitory neurons, but rather on the reach of their axonal arbor and the morphology of the post-synaptic neuron. This change of paradigm from pre-synaptic to post-synaptic view has very important consequences for the interpretation of the LFP in terms of activities of specific neuron types and the modelling of these signals. According to this paradigm, excitatory synapses on pyramidal neurons contribute little to the LFP, due to cancelling effects (Fig. 8A). However, their contribution can be still visible through di-synaptic mechanisms. In fact, the synapses of pyramidal neurons on basket cells are strong [33], so single action potentials of pyramidal neurons can activate reliably some basket cells. These, in turn, can produce IPSPs on pyramidal cells, which resulting LFP will thus be associated with the action potentials of the pyramidal neurons. Our model shows that this di-synaptic mechanism can lead to a measurable contribution of pyramidal neurons to the LFP, as observed experimentally. This also explains why in the presumed unitary field deduced from human recordings, inhibitory uLFP always have the same polarity and peak earlier than excitatory uLFP [44], consistent with the di-synaptic nature of the latter.

Thus, these findings help with the correct interpretation of the LFP signal. Not only the present modeling study provides a mechanistic explanation for previous experimental results, but it also suggests a new interpretation of the LFP signal. Because the LFP signal in the tissue is a sum of each neuron’s contribution (uLFPs), our paradigm predicts that the LFP mostly reflects the inhibitory currents in pyramidal cells. Note that this paradigm also predicts that soma-targeting interneurons should be much more visible in the LFP compared to dendrite-targeting inhibitory cells.

Can these considerations apply to more global signals recorded at the surface of the brain (electrocorticography, ECoG) or from the scalp (electroencephalography, EEG) ? Assuming that ECoG and EEG signals result from the electric dipoles made by pyramidal cells, the same considerations as above should apply. Our paradigm predicts that these signals should also be dominated by inhibitory activity, and reflect primarily the IPSPs on pyramidal cells in cortex. The testing of such a prediction should be investigated in future models.

Note that this interpretation assumes that the single-neuron contributions to LFP sum linearly, but in practice this summation may suffer from various non-linear effects. Deviations from a simple linear summation may result from different factors, such as the dense packing of dendritic processes in extracellular space, and the fact that there may be complicated spatial interactions between membrane and return currents. Extracellular conductivity may also be different in various regions of the neuropil, and extracellular space may also have diffusive and capacitive effects that may make the summation frequency dependent [4, 5]. These effects should be evaluated by more precise models taking these interactions into account.

Other limitations of the present study is that, first it did not include the LFP contribution of synaptic currents in inhibitory. However, these neurons are mostly spherically symmetric, so that their dipolar contribution is limited. Second, it did not include the possible contribution of intrinsic currents, which were shown to influence LFPs, such as Ih [36] or K+ conductances [14]. Including these currents in single-cell simulations showed moderate effects for excitatory LFPs and nearly no effect for inhibitory LFPs (Fig. 2), and the difference between the two was actually larger in the presence of intrinsic currents. Third, we did not consider the possible influence of glia which may also influence LFPs on a slow time-course through ionic buffering.

Finally, our approach provides a new way to calculate the LFP from networks of point neurons. Previously Hagen and colleagues [20] proposed calculating LFP generated from point neuron models by using their `hybridLFPy` set of Python classes. Their approach give good estimation of the field potential, however it requires a biophysical calculation of the field from a large number of neurons. Here, we propsoe an alternative approach which for the same type of models should give less precise but faster estimation of the field. In our model we calculate local field potential by convolving the unitary fields with the spiking activity of each point neuron type locating them in space. Those fields can be then summed linearly. The estimation of the field should be sufficient to estimate the LFP from networks of point neurons for better understanding of network activity (for example sharp waves in the hippocampus). This application will be developed in further work.

## Acknowledgments

This work was supported by Centre National de la Recherche Scientifique (CNRS, France), the European Community Future and Emerging Technologies program (The Human Brain Project, H2020-720270 and H2020-785907), the ANR PARADOX, and the ICODE excellence network. We would also like to thank Jose Donoso for valuable discussions.

